# Replacing non-biomedical concepts improves embedding of biomedical concepts

**DOI:** 10.1101/2024.07.01.601556

**Authors:** Enock Niyonkuru, Mauricio Soto Gomez, Elena Casiraghi, Stephan Antogiovanni, Hannah Blau, Justin T Reese, Giorgio Valentini, Peter N Robinson

## Abstract

**Objectives:** Concept embeddings are low-dimensional vector representations of concepts such as MeSH:D009203 (Myocardial Infarction), whose similarity in the embedded vector space reflects their semantic similarity. Here, we test the hypothesis that non-biomedical concept synonym replacement can improve the quality of biomedical concepts embeddings.

**Materials and methods:** We developed an approach that leverages WordNet to replace sets of synonyms with the most common representative of the synonym set.

**Results:** We tested our approach on 1055 concept sets and found that, on average, the mean intracluster distance was reduced by 8% in the vector-space. Assuming that homophily of related concepts in the vector space is desirable, our approach tends to improve the quality of embeddings.

**Discussion and Conclusion:** This pilot study shows that non-biomedical synonym replacement tends to improve the quality of embeddings of biomedical concepts using the Word2Vec algorithm. We have implemented our approach in a freely available Python package available at https://github.com/TheJacksonLaboratory/wn2vec.

## 1 Introduction

Word2vec is a two-layer neural network model that embeds individual words in a vector space, with the goal that similar words tend to be represented by close vectors (embeddings) in the vector space.^1^ The text to be embedded is represented by a corpus *W* of words *w* ∈ *W* and their contexts *c*. To compute the embeddings, the neural model is trained to predict the context words that surround the current word within a specific window. The goal is to find the network parameters *θ* maximizing the corpus probability; the optimal values for *θ* are the word embedding.^1, 2^ Numerous other effective word embedding models have been presented in the literature. ^3–6^ The original word2vec method operates on individual words (tokens); however, many biomedical concepts span multiple tokens. For instance, “bronchopulmonary dysplasia” would be treated by all the embedding models as two words representing two different semconceptsantics, while it represents a single medical concept (semantics). Therefore, recent “concept-replacement” approaches collapse multiword concepts into a single token or concept identifier^7^ (e.g., “Myocardial Infarction” is replaced by its MeSH id: D009203), so that the word-embedding algorithm treats the concept identifier (word) as a unique concept (related to a unique semantics). Existing approaches to concept replacement include the Narrative Information Linear Extraction (NILE) system to identify concepts from the Systematized Nomenclature of Medicine-Clinical Terms (SNOMED-CT) thesaurus^7–9^ and PubTator, which employs a series of specialized concept taggers to obtain annotations for each bioconcept type.^10, 11^

The central idea behind word embedding is that a word can be characterized by “the company it keeps”.^12^ That is the contexts in which a word appears to contain information about the meaning of the word. Therefore, concept replacement has the potential to improve the utility of word embeddings in two ways. First, concept replacement identifies multi-token concepts and replaces them with a (single-token) identifier as described above. Secondly, concept replacement replaces synonymous but distinct words and phrases with the same identifier (Myocardial Infarction, Myocardial Infarct, and Heart Attack would all be replaced by the same MeSH id D009203). Under the assumption that each of these synonyms would tend to have similar, yet varied contexts, synonym replacement would, on one hand, improve the informativeness of the embedding of a specific word by providing the embedding model with more varied contextual information about the word itself, i.e., more examples/contexts to learn from (i.e. all the contexts for all the synonyms of the word); on the other hand, by clustering all the examples from sets of synonym words, it would delete redundancies in the examples the word2vec network is provided with. A minor advantage is that the algorithm should have faster convergence due to the increased information and the smaller amount of words to be embedded. An empirical investigation showed that this improves the performance of medical-word emebedding.^7^

To our knowledge, however, previous efforts at concept replacement have been restricted to biomedical concepts. In this work, we reasoned that replacing synonyms of non-biomedical concepts with the same identifier would further improve the performance of the embedding. In this work, we present a simple heuristic for non-biomedical synonym replacement, we test it on a corpus containing 1,055 sets of related biomedical concepts and we show that our strategy embeds related biomedical concepts more close to each other than a purely biomedical concept replacement approach. In Figure 1 we sketch the pipeline of our experiments.

**Figure 1:**
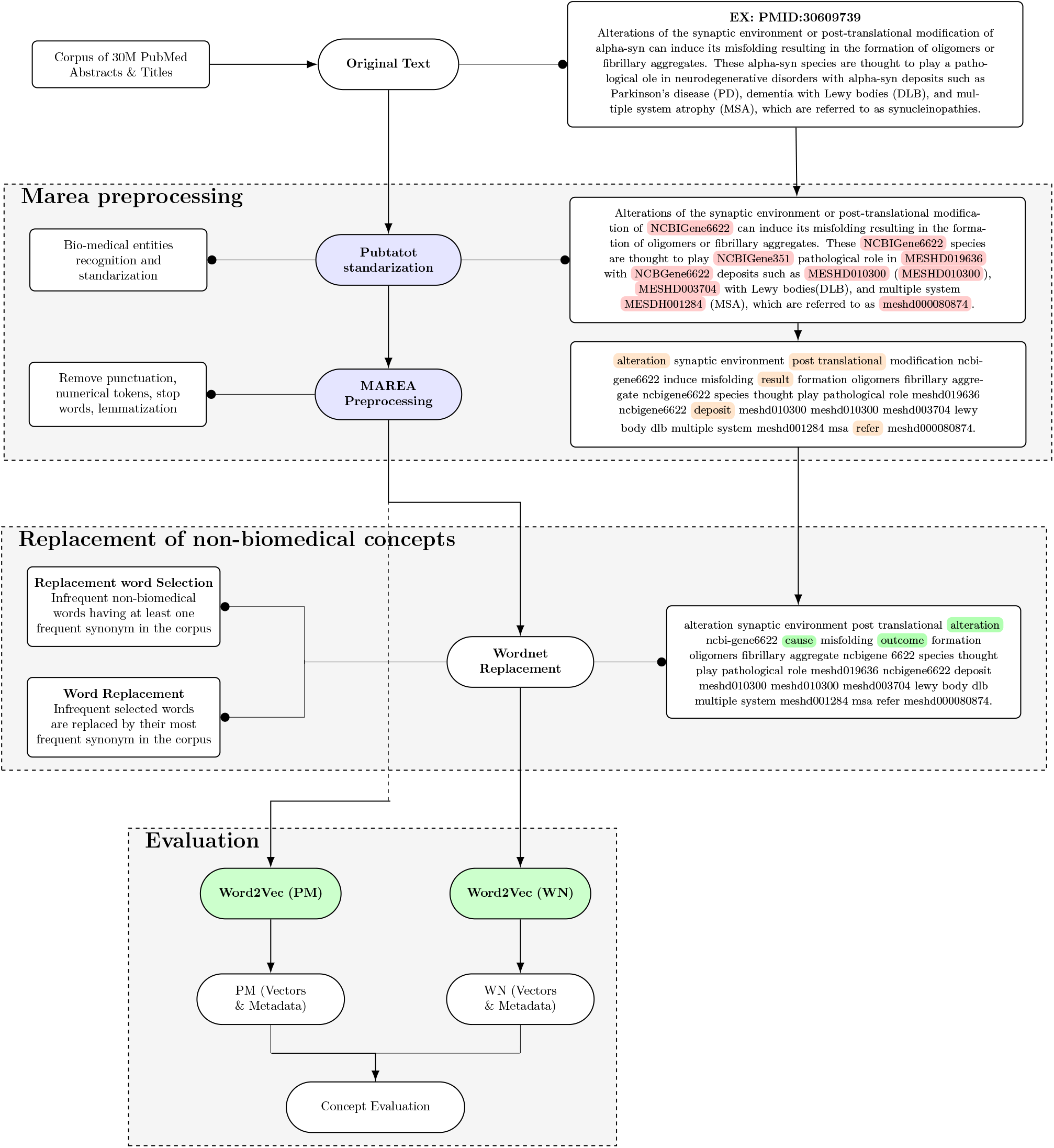
Schematic of the approach. The process starts with an initial text preprocessing, via Marea software, which allows obtaining the PM corpus. The PM corpus is further processed by our non-biomedical concepts replacement to obtain the WN corpus. Finally, word2vec embeds both the PM and the WN corpus, and the pairwise distances between sets of related biomedical concepts in the embedded PM corpus are compared to those in the embedded WN corpus to show the potential of our non-biomedical concept replacement strategy.

## 2. Materials and methods

### 2.1 Input corpus retrieval and text pre-processing with Marea

The corpus used to test our proposal is composed of 10,584,195 abstracts and titles published between January, 2010 and November, 2020 and available in PubMed; they were downloaded from the FTP site of the National Center for Biotechnology Information (NCBI) by using Marea, a software^13^1 that automatically parses the annual baseline and daily file-updates, p rovided i n t he f orm o f m etadata by N CBI, a nd extract the PubMed ID and year of publication of each paper.

Marea was also used to automatically pre-process the texts in the corpus, standardizing biomedical concepts and removing from unuseful information. In particular, Marea applies PubTator Central^10, 11^ concept recognition to standardize identifiers a nd h andle m ulti-word n oun p hrases t hat d esignate a single disease, chemical, or other entity. Following concept replacement, Marea eliminates punctuation, numerical tokens, and stop words, and reduces the vocabulary size through lemmatization.

### 2.2 Replacement of non-biomedical words by their WordNet synonym

The hypothesis of this research is that replacement of sets of highly related non-biomedical concepts by their common synonym will increase the ability of an embedding algorithm, e.g. word2vec, to place related biomedical concepts close to each other in vector space.

To identify synonyms of common words, we queried WordNet,^14^ a lexical database of English that groups nouns, verbs, adjectives, and adverbs into sets of cognitive synonyms (synsets), each expressing a distinct concept. Words are interlinked by conceptual-semantic and lexical relations (https://wordnet.princeton.edu/).

The replacement algorithm we devised starts by identifying the set of non-biomedical concepts (words) to be replaced. In particular, we reasoned that words frequently appearing in the corpus might be important and should not be replaced, together with infrequent words that have specific meanings (e.g. all their synonyms are also infrequent in the corpus) and might therefore carry detailed information. The other, infrequent and generic, words were replaced by their most frequent synonym in the corpus.

The set of words to be replaced, ℛ, is identified by computing the overall frequency, *f* (*w*), of each token, *w*, in the corpus (multiple occurrences in one abstract were counted multiple times).

Not-frequent words, i.e. words with *f* (*w*) *< τ*, being *τ* a user-set *replacement threshold* ^2^, are inserted in ℛ, and considered as candidates for replacement.

Then, ℛ is filtered to remove not-frequent words with specific meanings. To this aim, for each *w ∈* ℛ, we use WordNet to identify its synset, 𝒮_*w*_; next we select the synonym of *w* with the highest overall-frequency in the corpus, *s*_*max*_, and store it in a dictionary, **S**, mapping the word *w* to *s*_*max*_, i.e.

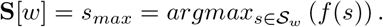

Words whose synonyms are all not-frequent in the corpus are easily recognized through the dictionary because *f* (*s*) *< τ, ∀s ∈* 𝒮_*w*_ *⇔ f* (**S**[*w*]) *< τ*. These words are considered as words with a specific, and possibly discriminatory meaning, and are therefore not replaced (removed from ℛ), while each other word *w ∈* ℛ is replaced by its most frequent synonym **S**[*w*].

The algorithm pseudo code is available in supplement as Algorithm 1, and a practical example of the replacement process is reported in Supplementary Figure S1.

### 2.3 Experiments

In the following sections we will refer to datasets processed only by Marea as PubMed-Marea, or “PM”; PM datasets further processed to substitute (not-frequent) non-biomedical concepts by their WordNet synonyms will be referred to as “WN”. While the number of unique biomedical concepts did not change between the PM and the WN corpus, the unique non-biomedical concepts in PM where larger then those in WN (3,018,918 in PM versus the 2,992,978 in WN)

We derived embeddings representing the concepts in the input corpora (the 10,579,997 PM or WN abstracts) by adapting the word2vec^1^ (word to vector embedding) implementation provided by the Gensim library.^15^ In particular, we used skipgram architecture with embedding size 128 (meaning that all concepts in PM and WN were represented as 128-dimensional vectors), window size 10, included words in the vocabulary that appear at least once in the corpus (mincount=1), and applied a sampling threshold of 10^*−*5^ for downsampling high-frequency words. The initial learning rate was set to 0.03 (alpha = 0.03) and was linearly decreased to a minimum of 0.0001 (min-alpha=0.0001) during training; we fixed the number of negative samples per positive context words to 5.

#### 2.3.1 Concept sets

Our assumption is that the quality of embeddings can be assessed by measuring the pairwise distances between the embeddings of related concepts.

To evaluate our proposal we therefore identified subsets of related genes and medical concepts prior to performing the testing and validation described in the following section ^3^.

To select gene subsets belonging to related pathways we analyzed the Molecular Signatures Database (MSigDB),^16^ from which we retrieved a total of 961 gene subsets (the number of gene subsets mined per data source is reported in Table 1). In addition, 94 subsets of related medical concepts were retrieved from Medical subject headings (MeSH) resource.^17^

**Table 1:** Comparison of mean interconcept distance for embedding with WordNet synonym replacement (WN) and without (PM). At different replacement thresholds, the columns WN *<* PM show the counts and percentages of concepts in each category for which WN embedding produced concept vectors that were closer to each other than PM, i.e., in which embeddings produced by our approach had a higher quality (shown in gray). The column PM *<* WN shows the opposite case. Data are shown for statistically significant (Sig) differences and for all comparisons (All). The “winner” in each comparison is shown in bold. Different thresholds (*τ*) were calculated based on the mean number of occurrences (427) of words considered for synonym replacement in our dataset.

| Replacement threshold ($\tau$) | Category | # sets | # Concepts | Significant | | All | |
| --- | --- | --- | --- | --- | --- | --- | --- |
|  |  |  |  | WN < PM | WN > PM | WN < PM | WN > PM |
| $0.5 \cdot \mu$ | MESH | 94 | 2503 | **12 (12.8%)** | 4 (4.3%) | 40 (42.6%) | **48 (51.1%)** |
|  | Biocarta | 285 | 1480 | **22 (7.7%)** | 19 (6.7%) | **145 (50.9%)** | 140 (49.1%) |
|  | KEGG | 182 | 4941 | 32 (17.6%) | 21 (11.5%) | 90 (49.5%) | **92 (50.5%)** |
|  | GO(bp) | 300 | 7030 | **36 (12.0%)** | 31 (10.3%) | **137 (45.7%)** | 135 (45.0%) |
|  | PID | 194 | 2507 | 18 (9.3%) | **26 (13.4%)** | 96 (49.5%) | **98 (50.5%)** |
| $1 \cdot \mu$ | MESH | 94 | 2503 | **13 (13.8%)** | 9 (9.6%) | **46 (48.9%)** | 42 (44.7%) |
|  | Biocarta | 285 | 1480 | **32 (11.2%)** | 12 (4.2%) | **154 (54.0%)** | 131 (46.0%) |
|  | KEGG | 182 | 4941 | **35 (19.2%)** | 13 (7.1%) | **112 (61.5%)** | 70 (38.5%) |
|  | GO(bp) | 300 | 7030 | **36 (12.0%)** | 27 (9.0%) | **147 (49.0%)** | 125 (41.7%) |
|  | PID | 194 | 2507 | **29 (14.9%)** | 5 (2.6%) | **115 (59.3%)** | 79 (40.7%) |
| $2 \cdot \mu$ | MESH | 94 | 2503 | **13 (13.8%)** | 6 (6.4%) | **49 (52.1%)** | 39 (41.5%) |
|  | Biocarta | 285 | 1480 | 22 (7.7%) | **28 (9.8%)** | **153 (53.7%)** | 132 (46.3%) |
|  | KEGG | 182 | 4941 | **37 (20.3%)** | 24 (13.2%) | **105 (57.7%)** | 77 (42.3%) |
|  | GO(bp) | 300 | 7030 | 34 (11.3%) | **37 (12.3%)** | **148 (49.3%)** | 124 (41.3%) |
|  | PID | 194 | 2507 | **27 (13.9%)** | 22 (11.3%) | **105 (54.1%)** | 89 (45.9%) |
| $4 \cdot \mu$ | MESH | 94 | 2503 | **14 (5.1%)** | 7 (2.5%) | **53 (19.2%)** | 35 (12.7%) |
|  | Biocarta | 285 | 1480 | 26 (8.8%) | **36 (12.1%)** | 121 (40.7%) | **164 (55.2%)** |
|  | KEGG | 182 | 4941 | **37 (11.6%)** | 21 (6.6%) | **96 (30.1%)** | 86 (27.0%) |
|  | GO(bp) | 300 | 7030 | **33 (9.7%)** | 32 (9.4%) | **144 (42.4%)** | 128 (37.6%) |
|  | PID | 194 | 2507 | 18 (5.0%) | **34 (9.4%)** | **89 (24.6%)** | 105 (29.0%) |

Concept subsets were deleted if they contained less than 5^4^ concepts that were represented in the test (PM or WN) corpus. For example, if a gene set had 100 genes but, in our corpus, only 3 genes belonging to gene set were present, then that gene set was deleted.

#### 2.3.2 Testing and Validation

We firstly checked that the scale and distribution of PM and WN vectorial space did not change. To this aim, we randomly sampled 1M vector-pairs in each dataset. We then calculated the distance between pairs of vectors and then plot the Empirical Cumulative Distribution Function (ECDFs) and Empirical Q-Q Plot of the computed distances (Figure 2). We visually verified that the two distributions had only slight differences.

**Figure 2:**
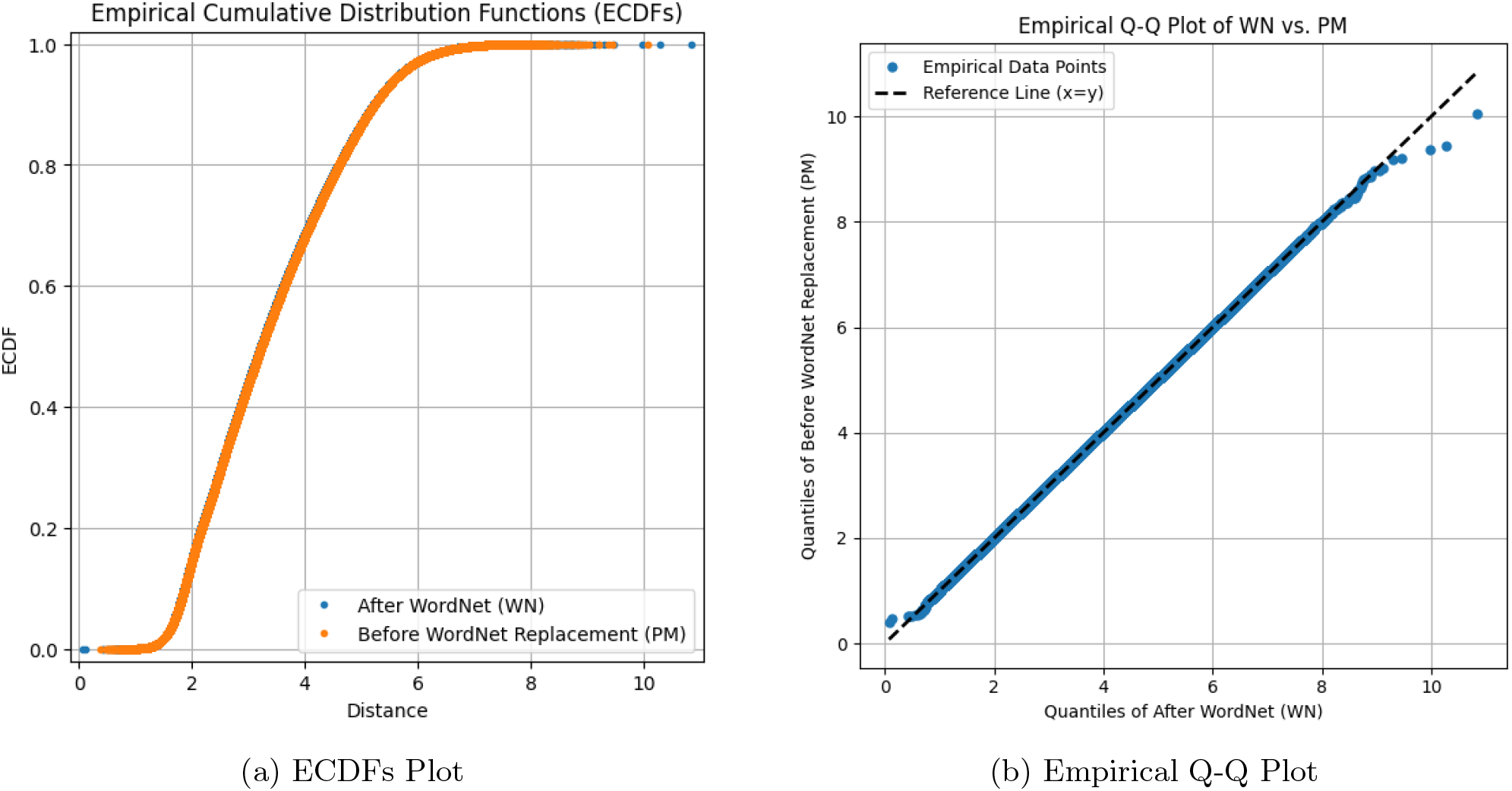
Comparative Analysis of WordNet Replacement Impact on Data Distribution. Figure (a) presents the Empirical Cumulative Distribution Functions (ECDFs), showcasing the cumulative frequency distribution before and after WordNet replacement, while Figure (b) illustrates the corresponding Empirical Q-Q Plot, detailing the quantile comparison between the original and the WordNet-replaced datasets. The close alignment of data points with the reference line in the Q-Q Plot and the overlap of the ECDF curves suggest minimal distributional deviation post-replacement.

Next, we analyzed the embedded representations obtained after PM and WN processing by focusing on individual subsets, *𝒳* (Section 2.3.1), and employing cosine similarity to evaluate all pairwise distances among the embedded concepts within *𝒳* ‘s representation post-PM processing versus post-WN processing. We then used the t-test to compare the pairwise-distances computed within the PM subset against those within the WN subset.

We observed that the application of our replacement strategy after Marea processing leads to an intracluster mean distance that is smaller than the one obtained when only Marea is applied. Indeed, over 1,055 sets of related gene and MeSH concepts sets, we found that, on average, the mean intra-cluster distance was reduced of the 8% - for sets where a significant difference was found, and by the 12% - on the average of all the comparisons (Figure 3).

**Figure 3:**
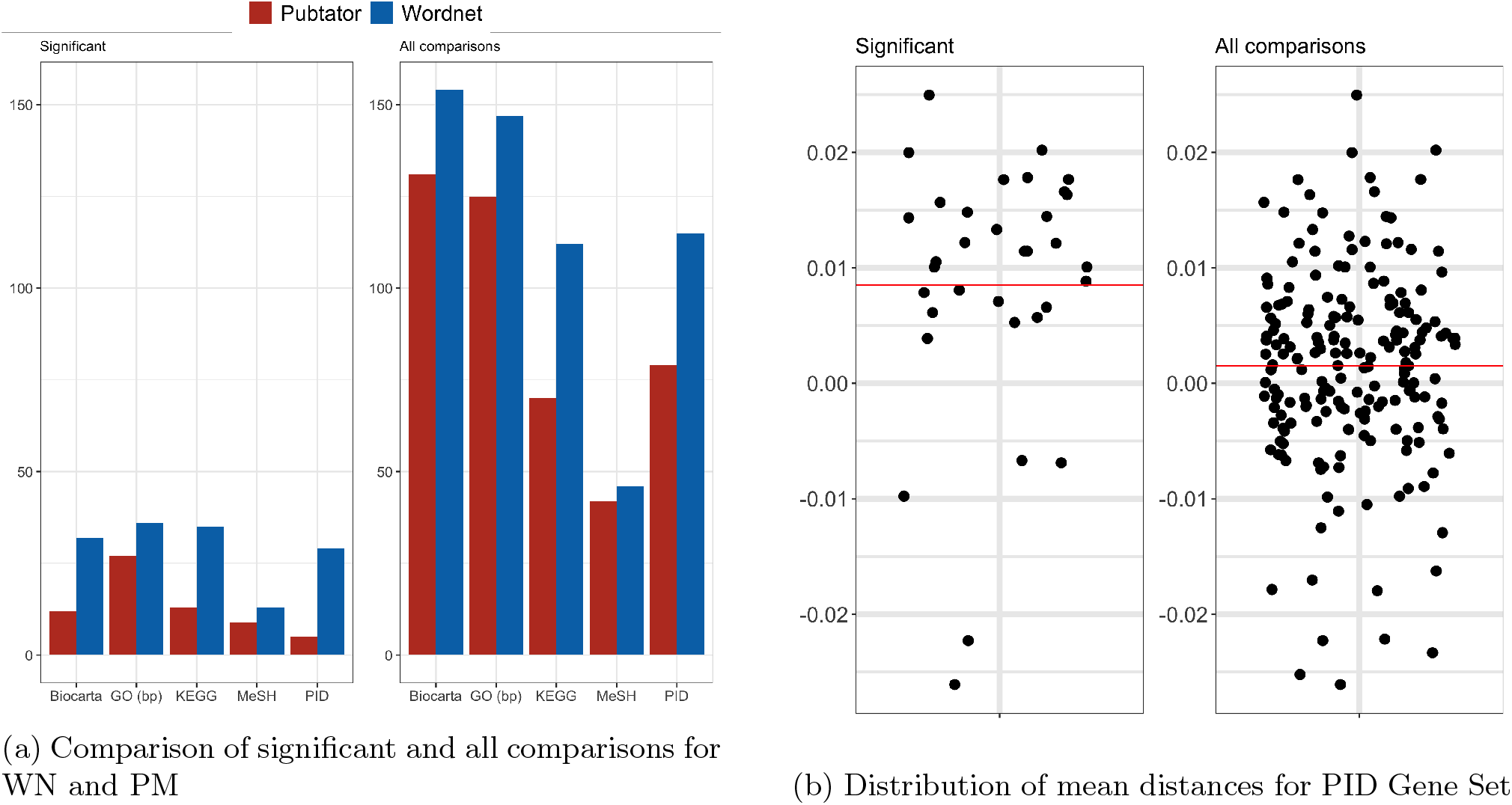
Comparative Analysis of WN and PM Methodologies: Figure (a) displays the bar chart comparing WN and PM across five distinct concept sets (Methods), highlighting the number of concept sets where the cluster mean distance is significantly lower, indicative of superior embeddings. ‘Significant’ designates those with a statistically significant difference in cluster mean distances (*p <* 0.05), while ‘All Comparisons’ encompasses the entire dataset. Figure (b) illustrates the spread of mean distances within the PID Gene Sets, detailing the variance and central tendency across 194 gene sets. ‘Significant’ encompasses gene sets with notable mean distance variations between ‘PM’ and ‘WN’ (*p <* 0.05), and ‘All comparisons’ includes all evaluated gene sets.

We also tested different thresholds for replacing non-biomedical terms (i.e., 214, 854, 1708, and the mean value of 427). We found that using the mean value yielded the best results (Table 1). Lower thresholds resulted in fewer words being replaced, while higher thresholds risked losing context by replacing too many words.

In addition, we investigated the impact of different parameters on the performance of our method. We varied the window size number, the higher the window size (i.e., 2, 5, and 10), the stronger the embeddings, and the more the WordNet synonym replacement had an impact on biomedical concept embeddings. The best results were obtained with a window size of 10 (Table 2).

**Table 2:** Comparison of window size for embedding with Wordnet synonym replacement (WN) and without (PM). The columns WN *<* PM show the counts and percentages of concepts in each category for which WN embedding produced concept vectors that were closer to each other than PM. The column PM *<* WN shows the opposite case. Data are shown for statistically significant (Sig) differences and for all comparisons (All). The “winner” in each comparison is shown in bold. This table describes the effect of window size on how the replacement of synonyms of non-biomedical concepts impacts the embedding of biomedical concepts. The analysis described in the text was performed at the window size of 10.0. Gray columns: same as in Table 1.

| Window Size | Category | # sets | # Concepts | Significant |  | All |  |
| --- | --- | --- | --- | --- | --- | --- | --- |
|  |  |  |  | WN < PM | WN > PM | WN < PM | WN > PM |
| 2 | MESH | 94 | 2503 | 9 (9.6%) | **10 (10.6%)** | 37 (39.4%) | **51 (54.3%)** |
|  | Biocarta | 285 | 1480 | **33 (11.6%)** | 30 (10.5%) | **152 (53.3%)** | 133 (46.7%) |
|  | KEGG | 182 | 4941 | **34 (18.7%)** | 20 (11.0%) | **101 (55.5%)** | 81 (44.5%) |
|  | GO(bp) | 300 | 7030 | 39 (13.0%) | **40 (13.3%)** | **144 (48.0%)** | 128 (42.7%) |
|  | PID | 194 | 2507 | **39 (20.1%)** | 24 (12.4%) | 96 (49.5%) | **98 (50.5%)** |
| 5 | MESH | 94 | 2503 | **14 (14.9%)** | 12 (12.8%) | 43 (45.7%) | **45 (47.9%)** |
|  | Biocarta | 285 | 1480 | **27 (9.5%)** | 21 (7.4%) | **152 (53.3%)** | 133 (46.7%) |
|  | KEGG | 182 | 4941 | **37 (20.3%)** | 17 (9.3%) | **104 (57.1%)** | 78 (42.9%) |
|  | GO(bp) | 300 | 7030 | **36 (12.0%)** | 27 (9.0%) | **144 (48.0%)** | 128 (42.7%) |
|  | PID | 194 | 2507 | **27 (13.9%)** | 20 (10.3%) | **102 (52.6%)** | 92 (47.4%) |
| 10 | MESH | 94 | 2503 | **13 (13.8%)** | 9 (9.6%) | **46 (48.9%)** | 42 (44.7%) |
|  | Biocarta | 285 | 1480 | **32 (11.2%)** | 12 (4.2%) | **154 (54.0%)** | 131 (46.0%) |
|  | KEGG | 182 | 4941 | **35 (19.2%)** | 13 (7.1%) | **112 (61.5%)** | 70 (38.5%) |
|  | GO(bp) | 300 | 7030 | **36 (12.0%)** | 27 (9.0%) | **147 (49.0%)** | 125 (41.7%) |
|  | PID | 194 | 2507 | **29 (14.9%)** | 5 (2.6%) | **115 (59.3%)** | 79 (40.7%) |

## 3 Discussion

Our pilot study demonstrates that replacing non-biomedical concepts tends to improve the homophily of word2vec-derived embeddings of related biomedical concepts, as assessed by the mean intra- and inter-distance between the embeddings of related and unrelated concepts. Although we focused on word2vec, other embedding algorithms could be as well used. Here, we employed a simple heuristic to perform non-biomedical synonym replacement, but more sophisticated approaches^18^ could further improve the embeddings. Code to implement our pipeline is available under an MIT license at https://github.com/TheJacksonLaboratory/wn2vec.

## Footnotes

1 Marea is freely available at https://github.com/TheJacksonLaboratory/marea.

2 In our experiments, *τ* = *µ*_*f*_, i.e. the mean of the overall frequency of all tokens in the corpus. This value was experimentally chosen (see Section 2.3.2)

3 The sets are available at the project GitHub site: https://github.com/TheJacksonLaboratory/wn2vec.

4 The minimum number of concepts in a set to be considered was fixed to 5 concepts under the assumption that larger sets would have less semantic focus.

