## Supplementary File for "Replacing non-biomedical concepts improves embedding of biomedical concepts"

**Algorithm 1** Non-Biomedical Word Replacement Algorithm

**Input:** Input corpus, WordNet, Replacement threshold $\tau$
**Output:** Modified corpus with replaced words

### S1 Selection of words to be replaced

Initialize set of words to be replaced $\mathcal{R} \leftarrow \emptyset$
Initialize set of high-frequency words $\mathcal{H} \leftarrow \emptyset$
Rank all tokens in the input corpus by their frequency
**for** each word $w$ in the corpus **do**
    **if** $f(w) < \tau$ **then**
        Add $w$ to $\mathcal{R}$
    **else**
        Add $w$ to $\mathcal{H}$
    **end if**
**end for**
Initialize dictionary $\mathbf{S} \leftarrow \emptyset$
**for** each word $w \in \mathcal{R}$ **do**
    **if** $\mathcal{S}_w \cap \mathcal{H} = \emptyset$ **then**
        Remove $w$ from $\mathcal{R}$
    **else**
        $\mathcal{S}_w \leftarrow$ WordNet synset of $w$
        $\mathbf{S}[w] \leftarrow \arg\max_s \{f(s), s \in \mathcal{S}_w\}$
    **end if**
**end for**

### S2 Replacement process

**for** each word $w \in \mathcal{R}$ **do**
    Replace $w$ in the corpus with $\mathbf{S}[w]$
**end for**

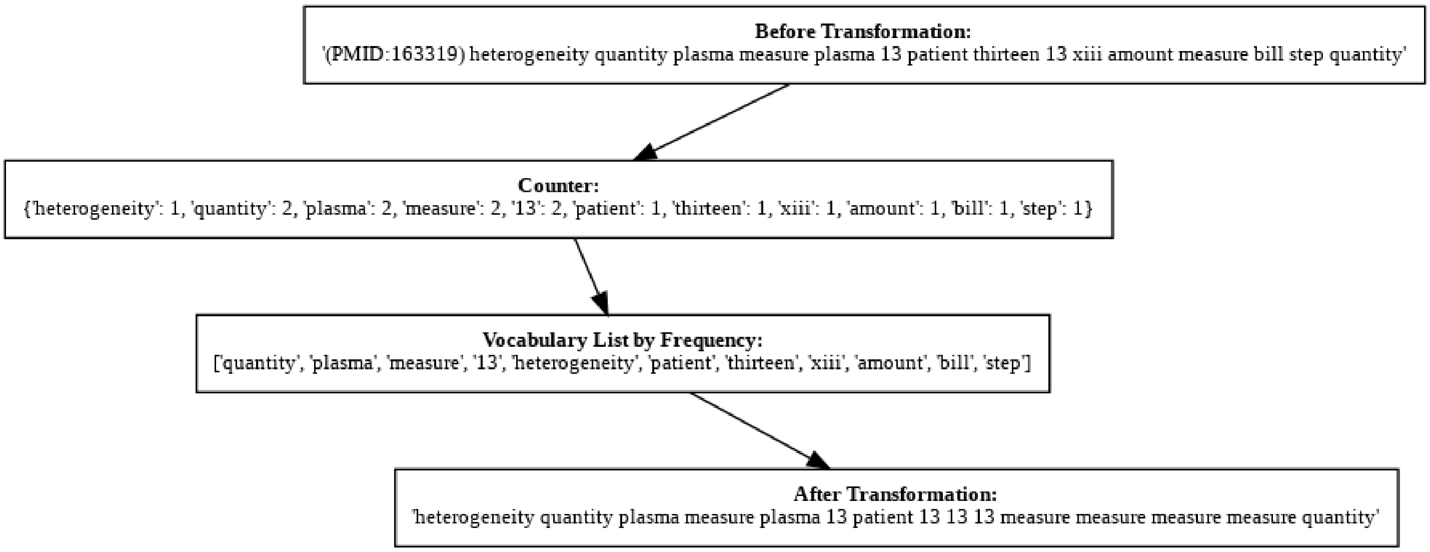


**Figure S1. Illustration of the Text Transformation Process** . The process starts with an initial text segment ('Before Transformation'), followed by a word frequency count ('Counter'). This is used to generate a 'Vocabulary List by Frequency', which then informs the final modified text ('After Transformation'). The procedure exemplifies a methodical approach of the algorithm to replace non-biomedical tokens.
